# A Comprehensive Reference Catalog of Human Skin DNA Virome Reveals Novel Viral Diversity and Microenvironmental Influences

**DOI:** 10.1101/2025.03.20.644480

**Authors:** Zhiming Li, Shenghui Li, Chongyin Han, Yuxiao Chen, Hefu Zhen, Yuzhe Sun, Xiaofeng Zhou, Yanmei Chen, Yan Zheng, Lianyi Han, Jean Krutmann, Chao Nie, Jiucun Wang, Jingjing Xia

**Affiliations:** State Key Laboratory of Genetic Engineering, Collaborative Innovation Center for Genetics and Development, School of Life Sciences, Fudan University, Shanghai, China; BGI Research, Shenzhen 518083, China; Shenzhen Key Laboratory of Neurogenomics, BGI Research, Shenzhen 518083, China; Key Laboratory of Precision Nutrition and Food Quality, Department of Nutrition and Health, China Agricultural University, Beijing, China; Greater Bay Area Institute of Precision Medicine (Guangzhou), School of Life Sciences, Fudan University, China; Department of Dermatology, Huashan Hospital, Fudan University, Shanghai, China; IUF Leibniz Research Institute for Environmental Medicine, Düsseldorf, Germany; Human Phenome Institute, Fudan University, Shanghai, China

**Keywords:** Skin virome, Metagenomic, Skin microenvironments, virusal catalog, Skin phages

## Abstract

**Introduction:** Human skin serves as a dynamic habitat for a diverse microbiome, including a complex array of viruses whose diversity and roles are not fully understood.

**Objectives:** This study aims to enhance our understanding of this skin viral diversity through the construction of a detailed, non-redundant DNA viral reference catalog.

**Methods:** A total of 2,760 skin metagenomes from six published skin studies were collected. A skin virome catalog was constructed using methods independent of reference databases. Viral characteristics were identified through cross-cohort meta-analysis and used to characterize viral features across different skin environments.

**Results:** We identified 20,927 viral sequences, which clustered into 2,873 viral operational taxonomic units (vOTUs). Among these, 2,610 represent previously unrecorded viral sequences, uncovering a substantial breadth of viral diversity on human skin. The results also highlight significant differences in viral communities that are associated with varying skin microenvironments. The oily skin is enriched in *Papillomaviridae*; the dry skin area is enriched in *Autographiviridae*, *Inoviridae* and *Mitoviridae*; and the moist skin is enriched in *Herelleviridae*, indicating the adaptive nature of viruses. We also investigated the relationship between bacteriophages and bacteria on the skin surface. We found that skin bacteria such as *Pseudomonas*, *Klebsiella*, and *Staphylococcus* are infected by phages from the class *Caudoviricetes*.

**Conclusion:** This comprehensive skin DNA viral catalog significantly advances our understanding of the virome’s role within the skin ecosystem. The findings highlight the adaptive nature of viruses to different skin microenvironments and their interactions with resident bacteria. This catalog serves as a valuable resource for further epidemiological and therapeutic research, potentially leading to better management and treatment of skin conditions influenced by the skin’s virome.

## Introduction

The skin, the largest organ of the human body, not only acts as a protective barrier against chemical injury, ultraviolet radiation and pathogens, but also harbours a complex ecosystem of microorganisms including bacteria, fungi, archaea and viruses^1,2^. This microbial community, known collectively as the skin microbiome, plays a crucial role in modulating immune responses, protecting against pathogen colonization and maintaining skin health^3, 4^. While considerable research has focused on the bacterial or fungal component of the skin microbiome, the viral component, its diversity and function, remains less explored. Existing research indicates that viruses account for approximately 1.51% of the microbial population on facial skin samples^5^. There is also an observed phenomenon known as “viral bloom” in specific samples, where viruses can comprise up to 96% of the microbial community at certain skin sites^6^. The most prevalent viruses on the skin surface are bacteriophages, including those that target *Cutibacterium* and *Staphylococcus*. Additionally, potential human viral pathogens such as human papillomavirus and Merkel cell polyomavirus are also present^6^.

Metagenomic sequencing and bioinformatics have provided tools to delve deeper into the virome’s complexity, without the limitations of traditional culture-based techniques. Viruses, particularly bacteriophages, which are viruses that infect bacteria, are now recognized as pivotal players in shaping microbial community dynamics. They can influence bacterial diversity and density through lytic cycles—where viruses replicate within and lyse their bacterial hosts—and lysogenic cycles—where viral genomes integrate into the bacterial genome, potentially altering host bacterial behavior and pathogenicity^7-9^.

The exploration of the skin virome is crucial for several reasons. First, viruses can influence the health and disease states of the skin by impacting microbial balance and host immune responses^10-12^. For example, bacteriophages can control the population of pathogenic bacteria, thereby preventing or exacerbating skin conditions such as acne, atopic dermatitis, and wound infections^13, 14^. Second, understanding the skin virome might open up therapeutic avenues, such as phage therapy, to combat antibiotic-resistant bacterial strains^14^. Finally, the skin virome can serve as a diagnostic marker for certain skin diseases and potentially for systemic disorders, given the skin’s interaction with other organs^15-17^.

To enhance our understanding of the skin virome, we have developed a non-redundant viral reference catalog based on human skin samples collected worldwide. This project involved the analysis of 2,760 metagenomic datasets from the United States, China, Singapore, and Italy, covering a variety of microenvironmental conditions. Our goal in establishing this catalog was to map the diversity of the skin virome, investigate its interaction with host and environmental factors, and assess its potential roles in skin health and disease.

## Material and methods

### Data collection

We conducted a comprehensive review of studies focused on skin microbiomes, excluding those that relied solely on ITS or 16S sequencing due to their limitations in capturing viral genomic data, which is essential for our cataloging objectives. Our inclusion criteria were limited to studies utilizing metagenomic sequencing techniques without enrichment for virus-like particles. We did not filter studies based on variations in sequencing platforms, extraction methods, or library preparation techniques. We collected and downloaded a total of 2,760 skin metagenomic data sets from the NCBI Sequence Read Archive (SRA) databases at (https://www.ncbi.nlm.nih.gov/sra) and CNGB (https://db.cngb.org/). To provide a clear demographic context for our data, we summarized the age, gender, country, and specific skin sites of the subjects in Table S1, with additional metadata detailed in Table S2. In the collected data sets, 97.72% (2,697) are generated from healthy individuals, 1.41% (39) are from persons with atopic dermatitis, and 0.87% (24) are from persons with psoriasis.

### Data processing

The raw reads were filtered using SOAPnuke (v2.1.9, with default parameters)^18, 19^. Human reads were further removed by aligning the filtered reads with the human genome hg38 using bowtie2 (v2.3.5.1, with default parameters)^20^. The remaining reads of each data set were assembled *de novo* using MEGAHIT (v1.2.9)^21^ based on various k-mer sizes (k=21, 33, 55, 77, 99).

### Virus Identification and Decontamination

Contigs larger than 2000 nt were preliminarily screened through VIBRANT (v1.2.1)^22^and DeepVirFinder (v1.0)^23^ (p-value <0.01 and score >0.90), identifying 2,480,992 contigs as potential viral sequences. To purify these viral sequences, following previous studies^24^, we utilized hmmsearch (v3.1)^25^ with default options to search for bacterial, archaeal and fungal universal single-copy orthologs (BUSCO-v5.4.4)^26^ within the viral sequences and removed them. Subsequently, we used BLASTN to align the resulting viral sequences against the human genome to remove sequences potentially originating from human genomic regions. Finally, we used geNomad^27^ (v1.8.1, with default parameters) to further filter the sequences (virus score of at least 0.7) and employed CheckV^28^ (v1.0.3, with default parameters) to assess the quality and completeness of single-contig virus genomes. Through these steps, we identified 4,199 complete viral genomes, 5,249 high-quality viral genomes with >90% completeness, and 11,479 medium-quality viral genomes with 50-90% completeness.The contigs were clustered through the following steps: (1) Use BLASTN alignment to calculate the similarity and coverage of contigs. (2) The HipMCL algorithm^29^ was used to perform sequence clustering based on 95% similarity and 85% alignment coverage, resulting in 2,873 potential viral operational taxonomic units (vOTUs), yielding 558 complete viruses, 841 high-quality viruses, and 1,474 medium-quality viruses.

### Viral taxonomy

We annotated vOTUs based on the annotation results from geNomad^27^(v1.8.1, with default parameters, This classification method follows the taxonomy in ICTV’s VMR number 19). In the SVD catalog, 98.75% (2,837 out of 2,873) of the viral operational taxonomic units (vOTUs) were classified.

### Host prediction

Host prediction for vOTUs was carried out using iPHoP^30^ (version 1.3.3, default parameters). The database was constructed using a published collection of skin microbiome genomes^31^. The construction process strictly followed the procedure outlined in iPHoP under “Adding bacterial and/or archaeal MAGs to the host database”.

### Functional annotation

In the Skin Virome Database (SVD), protein-coding sequences of viral operational taxonomic units (vOTUs) were predicted and annotated using Prodigal and EggNOG (v 5.0)^32^.

### Phylogenetic analysis

Phylogenetic analysis was conducted on 2,873 vOTUs from the Skin Virome Database (SVD) using a phylogenetic approach based on amino acid sequence similarity. We utilized ViPTreeGen (v1.1.3)^33^ to generate a viral proteomic tree of the SVD. The proteomic tree was then visualized using iTOL (v6.9)^34^.

### Quantification of vOTUs

To explore the biogeographical characteristics of the skin virome, we analyzed metagenomic data collected from oily, moist, and dry skin areas, aligning it against all vOTUs in the Skin Virome Database (SVD), covering a total of 2,334 metagenomic datasets. We used bowtie2 with a□>□95% identity threshold to align high-quality reads with all vOTUs in the SVD. Subsequently, we conducted a sequence-based vOTU abundance analysis using jgi_summarize_bam_contig_depth (default parameters)^35^. The mapping of reads to the SVD generated counts, forming a coverage depth/abundance matrix. Considering that different samples might have varying sequencing depths, we used a normalized coverage depth matrix to estimate the abundance of vOTUs. This approach allows us to fairly compare the relative abundance of vOTUs across different samples, thereby revealing the variations in viral composition across different skin types.

### Phage-host association analysis

To explore the associations between phage vOTUs and hosts within the community, we assessed the Spearman correlation between the relative abundances of phage vOTUs and microbial species. If a phage vOTU is predicted to infect a single genus (i.e., a specialist phage), we calculate the correlation between that phage vOTU and its predicted host. If a phage vOTU is predicted to infect multiple genera (i.e., a generalist phage), we calculate the correlation for each predicted host. If a phage-host pair (with zero abundance) does not exist in the sample, that sample is excluded from the correlation analysis.

### Alpha diversity

Alpha diversity is estimated based on the relative abundances of phage vOTUs. The Shannon diversity index is calculated by calling the *diversity* function in R (V4.3.1) using the *vegan* package^36^, with the parameter set to index = shannon. Comparisons of alpha diversity across different microenvironments and sites are conducted using the Wilcoxon rank-sum test, and a p-value of less than 0.05 indicates a significant difference.

### Statistical analysis

Using the ade4 package^37^ within the R platform, PCoA and PCA is performed on samples from different microenvironments and sites. Permutational Multivariate Analysis of Variance (PERMANOVA) is performed using the adonis function from the vegan package. When using PERMANOVA, the skin microenvironment and skin site characteristics is analyzed after adjusting for age and gender. Statistical significance is verified using the wilcox.test and kruskal.test functions. P-values are adjusted using the p.adjust function with the parameter method = fdr. Adjusted P-values less than 0.05 are considered to be statistically significant.

### Data availability

The data files of SVD, including nonredundant viral genomes, annotations, gene sequences and protein sequences, have been deposited in the https://github.com/lizhiming11/skin_virus. For detailed information on the metagenomic data, please refer to Supplementary Table 1.

### Code availability

The data preprocessing, genomic, functional analysis and statistical scripts of this study are available at https://github.com/lizhiming11/skin_virus.

### Compliance with Ethics Requirements

All procedures followed were in accordance with the ethical standards of the BGI Review Board of Bioethics and Biosafety (BGI-IRB21165). Informed consent was obtained from all samples for being included in the study.

## Results

### The construction of the Human Skin Viral Catalog

To this end, we collected a total of 2,760 publicly available metagenomic data from human skin, establishing a skin metagenomic dataset that covers the United States, China, Singapore, and Italy (Table S1). This dataset comprises samples from 21 different skin sites, categorized into three skin microenvironments: oily (1,849 samples), dry (244 samples), and moist (241 samples). After uniform processing (reads preprocessing and host removal), these samples represented 34.11 Tb of high-quality non-human metagenomic data and were used to generate a total of 250.56 million long contigs (Table S2). Using an integrated pipeline based on homology and features (see Methods), approximately 0.96% (n = 2,422,445) of the contigs were detected as highly credible viral sequences (Figure S1). The results showed that there were 4,199 complete viral genomes, 5,249 high-quality viral genomes, and 11,479 medium-quality viral genomes (Figure S1 and TableS3). The viral sequences were clustered according to MIUViG standards, with >95% nucleotide similarity and 85% alignment coverage^38^, forming a skin virome database (SVD) consisting of 2,873 viral operational taxonomic units (vOTUs) (Figure S1). This database included 558 complete viral genomes, 841 high-quality viral genomes, and 1,474 medium-quality viral genomes (Figure 1a and Table S4). The length of the vOTUs ranged from 2,010 bp to 433,960 bp, with an median length of 28,261 bp (Table S4).

**Figure 1.**
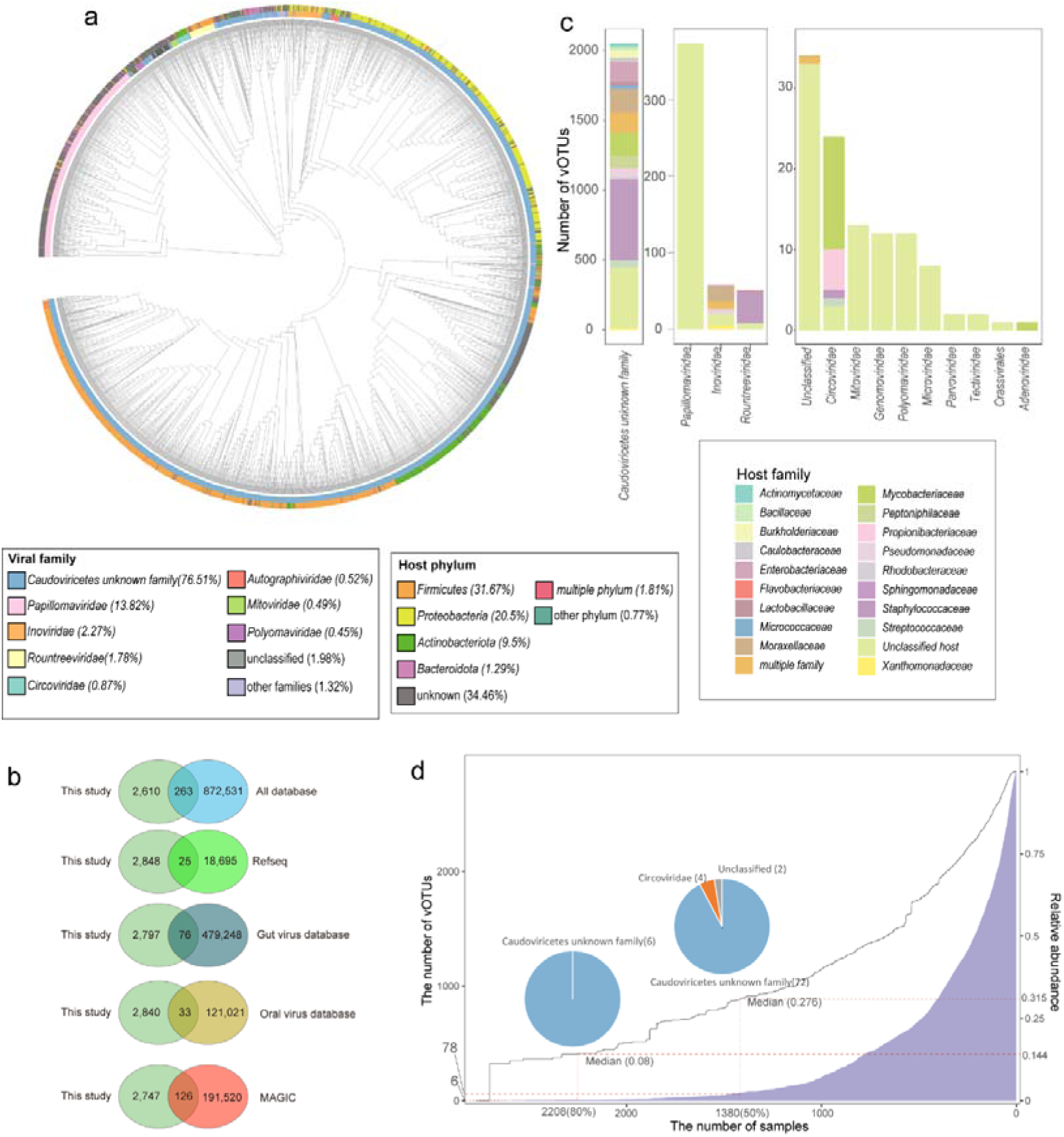
Overview of Viral Operational Taxonomic Units (vOTUs) of skin. **a.** Proteomic tree of 18,241 high-quality vOTUs. This tree was generated using ViPTreeGen. The outer ring displays the host classification for each vOTU: the innermost circle represents the viral family-level classification assignments. **b.** Comparison of vOTUs identified in this study with those in databases. The Venn diagram illustrates the overlap between vOTUs identified in this study and those in public databases. The clustering of phage genomes has a threshold of >95% identity and >85% length coverage. **c.** The bar chart displays the classification of the top 14 skin vOTUs. The host classification groups of the vOTUs are shown in different colors. **d**. The ubiquity of skin surface viruses in the samples is depicted through multiple graphical representations. The blue bar chart (corresponding to the left Y-axis) shows the number of viral Operational Taxonomic Units (vOTUs) that appear in a given number of samples (X-axis). The dashed lines indicate the quantity of vOTUs and their corresponding total relative abundance for sample sizes of 2208 and 1380. Pie charts display the proportion of vOTUs corresponding to each family when the sample size is 2208 and 1380. A linear curve (corresponding to the right Y-axis) quantifies the cumulative relative abundance of these vOTUs in the samples.

The vOTUs of the SVD were compared with several existing viral catalogs, including five human gut virome catalogs^24, 39-42^, two human-associated comprehensive viral catalogs (Cenote Human Virome Database (CHVD)^43^ and virushostdb^44^), one oral virome database(OVD)^45^, a Metagenome-Assembled Genome Inventory for Children (MAGIC)^46^ and available viral genomes in the RefSeq database. Of these, using a 95% similarity and 85% coverage threshold, only 263 vOTUs in the SVD matched with viral sequences in known databases, whereas 2,610 vOTUs did not match any known genomes in the databases (Figure 1b and Figure S2), indicating that they were previously unrecognized vOTUs.

The 2,873 vOTUs encoded 119,826 protein-coding genes collectively. Using the eggNOG database, a comprehensive resource for classifying genes into orthologous groups^32^, it was found that 70,821 (59.1%) of these genes were not annotated in the eggNOG database, and 28,447 (23.74%) had unknown functions in the eggNOG database (Figure S3a). The principal coordinate analysis (PCoA) of the vOTUs functional profiles revealed differences in the spatial distribution of vOTUs. (Figure S3b, PERMANOVA test, p < 0.01). Different viral families show variations in functions related to DNA repair, DNA replication, peptidases and inhibitors, and replication and repair processes (Figure S4). For example, the *Autographiviridae* are functionally enriched in DNA repair and recombination proteins, while the *Herelleviridae* were functionally enriched in replication and repair, and peptidases and inhibitors (Figure S4). This variety may reflect the adaptation strategies of viruses to their host environments. For example, variations in functions related to DNA repair and replication indicate how different viruses respond to host cellular defense mechanisms^47^. Such adaptability is a key driver of viral evolution and has significant implications for the infection strategies of the viruses and their survival capabilities.

### Taxonomic landscape and host range of skin viruses

The taxonomic annotation results from geNomad^27^ were utilized in the SVD catalog, with 98.02% (2,816 out of 2,873) of the vOTUs being classified. Notably, the majority, 76.51% (2,198), were assigned to unidentified taxonomic categories within the class *Caudoviricetes*. The viral families identified within the SVD include *Papillomaviridae* (n=397), *Inoviridae* (n=65) and *Rountreeviridae* (n=51) (Figure 1ac). *Papillomaviruses* were a group of ancient viruses that had largely co-evolved with their hosts, primarily colonizing the human skin surface and hair follicles^48^.

We found that at least 6 vOTUs, which were belonged to the class *Caudoviricetes*, appeared in at least 2,208 samples (representing 80%) and accounted for 14.4% (median 8%) of the total viral abundance (Figure 1d). In at least 1,380 (50%) of the samples, 78 vOTUs were identified, representing 31.5% (median 27.6%) of the total abundance, primarily including 72 members of unidentified families within the order *Caudoviricetes*, 2 from *Circoviridae*, and 2 Unclassified vOTUs (Figure 1d). Those commensal viral species might play a conserved role in human skin.

We used iPHoP to predict skin bacteria that would be infected by vOTUs. Among 1,880 phages (79.53%) were predicted to belong to vOTUs capable of infecting bacterial species on the skin surface. (Table S4). Among the 57 vOTUs that were not classified, 3 were predicted to infect skin bacteria (Table S4). The predicted infected bacteria include members of the phylum *Firmicutes* (n=910), followed by *Proteobacteria* (n=589), and *Actinobacteriota* (n=273) (Figure S5ab and Table S4). At the family level, the bacteria infected include *Staphylococcaceae* (n = 603), *Moraxellaceae* (n = 191), *Mycobacteriaceae* (n = 164), *Enterobacteriaceae* (n = 147), and *Peptoniphilaceae* (n = 92) (Figure S5c). At the genus level, the predicted infected skin bacteria include *Staphylococcus* (n = 601), *Corynebacterium* (n = 149), *Acinetobacter* (n = 94), *Moraxella* (n = 66), and *Anaerococcus* (n = 51) (Figure S5d). In addition to these abundant taxa, the phages were also predicted to infect *Enterococcus* (n = 15), *Klebsiella* (n = 76) and *Pseudomonas* (n = 56) (Table S4). Among all the phages predicted to infect skin bacteria (n=1956), the majority (78.81%, 1484) infect only a single genus (i.e., specialist bacteriophages vOTUs), whereas the others (21.19%, 399) infect multiple genera (e.g., generalist bacteriophages vOTUs) (Figure S5d).

Further analysis revealed that unclassified phages within *Caudoviricetes* can infect major bacteria in the human skin microbiome, including members of the *Propionibacteriaceae*, *Mycobacteriaceae*, *Staphylococcaceae*, *Peptoniphilaceae* and *Moraxellaceae* (Figure 2a). *Caudoviricetes* phages belonging to specific families such as *Autographiviridae, Herelleviridae, Rountreeviridae, Crassvirales,* and *Schitoviridae* are capable of infecting *Pseudomonadaceae, Caulobacteraceae, Enterobacteriaceae, Staphylococcaceae, Mycobacteriaceae,* and *Streptococcaceae* (Figure 2a). Among the unclassified phages from the class *Caudoviricetes*, specialist phages can infect common skin bacteria such as *Staphylococcus* (n = 548), *Corynebacterium* (n = 143), *Acinetobacter* (n = 73), and *Moraxella* (n = 66) (Figure 2b). Within the *Rountreeviridae* family of *Caudoviricetes*, specialist phages infect *Staphylococcus* (n=42) and *Streptococcus* (n=1). (Figure 2c). In the family of *Inoviridae*, specialist phages infect *Acinetobacter* (n=17), *Neisseria* (n=6), *Pseudomonas* (n=2), and *Xanthomonas* (n=2) (Figure 2c). Among the generalist phages, unclassified vOTUs from the class *Caudoviricetes* were found to infect *Klebsiella* and *Enterobacter* as well as *Staphylococcus* and *Acinetobacter*, all of which include multidrug-resistant strains ^1^. This implied the potential of *Caudoviricetes* phages in combating multidrug-resistant bacteria (Figure 2d).

**Figure 2.**
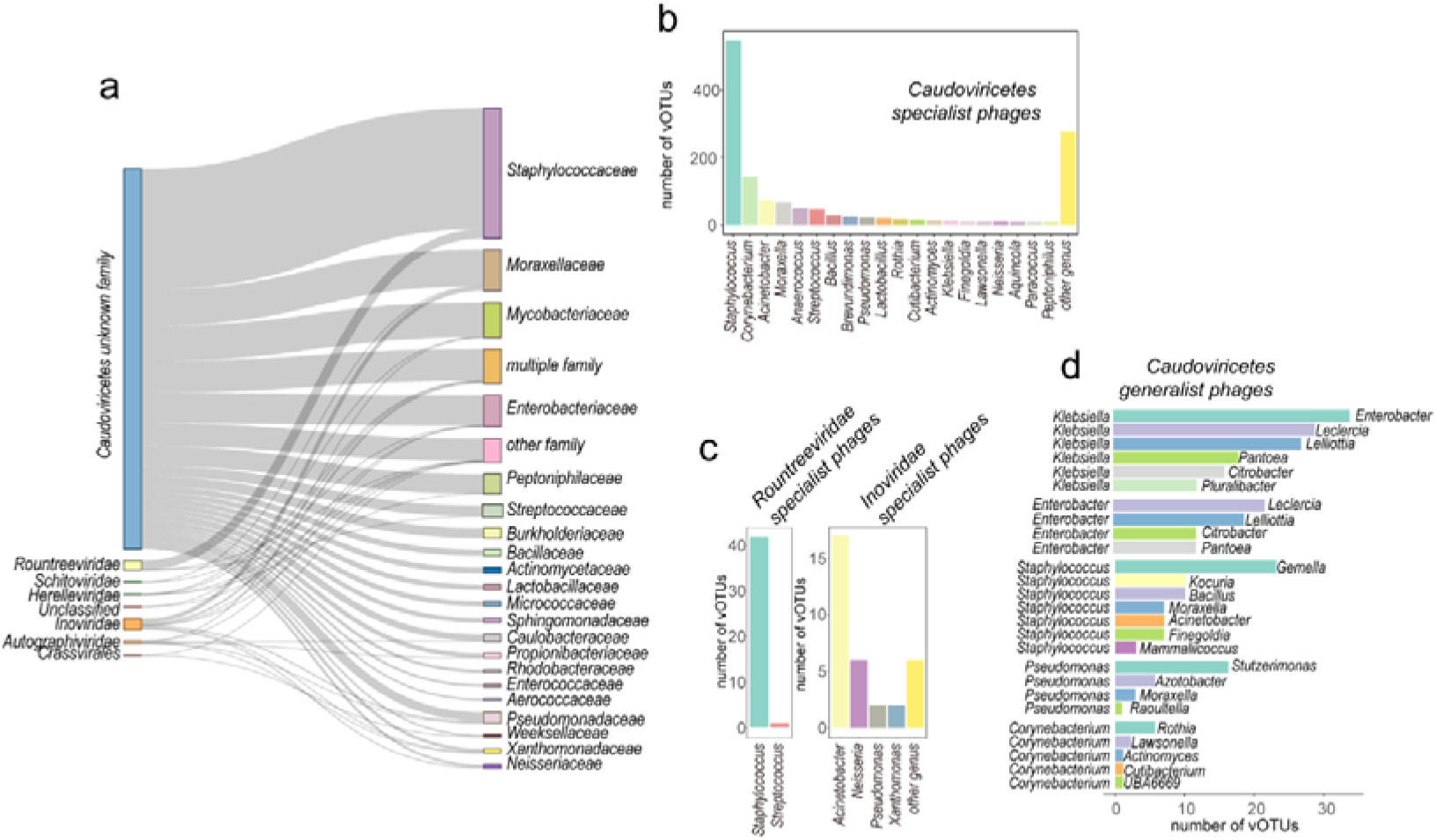
General overview of bacteriophages vOTUs on the skin surface. **a.** A Sankey diagram depicting skin vOTU at the family level and their corresponding hosts determined by iPHoP matching. The width of the links represents the number of relationships, with wider links indicating a greater number. **b-c.** The distribution of specialist phages vOTUs within the *Caudoviricetes* (b), *Rountreeviridae* and *Inoviridae* (c), showing the specific hosts they infect. **d.** The distribution of hosts for generalist phages vOTUs within the *Caudoviricetes*. The numbers indicate the count of vOTUs that infect two specific bacterial hosts.

### Close interactions between the skin phages and bacteriome

To explore the interaction between phages and bacteria in the skin, phages and bacterial profiles collected from 2,760 samples were compared. Results showed that the α-diversity (Shannon diversity) and β-diversity (Bray-Curtis distance) were significantly positively correlated (rs = 0.607 and 0.645, respectively; Figure 3ab), suggesting a close phages-bacteriome interaction.

We then analyzed the one-to-one correlations between the relative abundances of phages and their predicted host at the genus level across 2,760 samples. We found a positive correlation with an average rs of 0.357 between them (Figure 3c), consistent with previous findings in the gut (rs = 0.18) that phages and their host bacterial species coexist rather than exclude each other in human skin^49^. Additionally, specialist phages phages (average rs = 0.359) exhibited a higher correlation with their hosts compared to generalist phages vOTUs (average rs = 0.341) (Figure 3c). Upon inspected the relationships between each genus-level bacteria and their corresponding phages vOTUs; we found that, except for *Corynebacterium* and *Staphylococcus,* the relationships were greater for specialist phages vOTUs than for generalist phages, including common skin genera such as *Moraxella*, *Pseudomonas*, and *Streptococcus*(Figure 3d).

**Figure 3.**
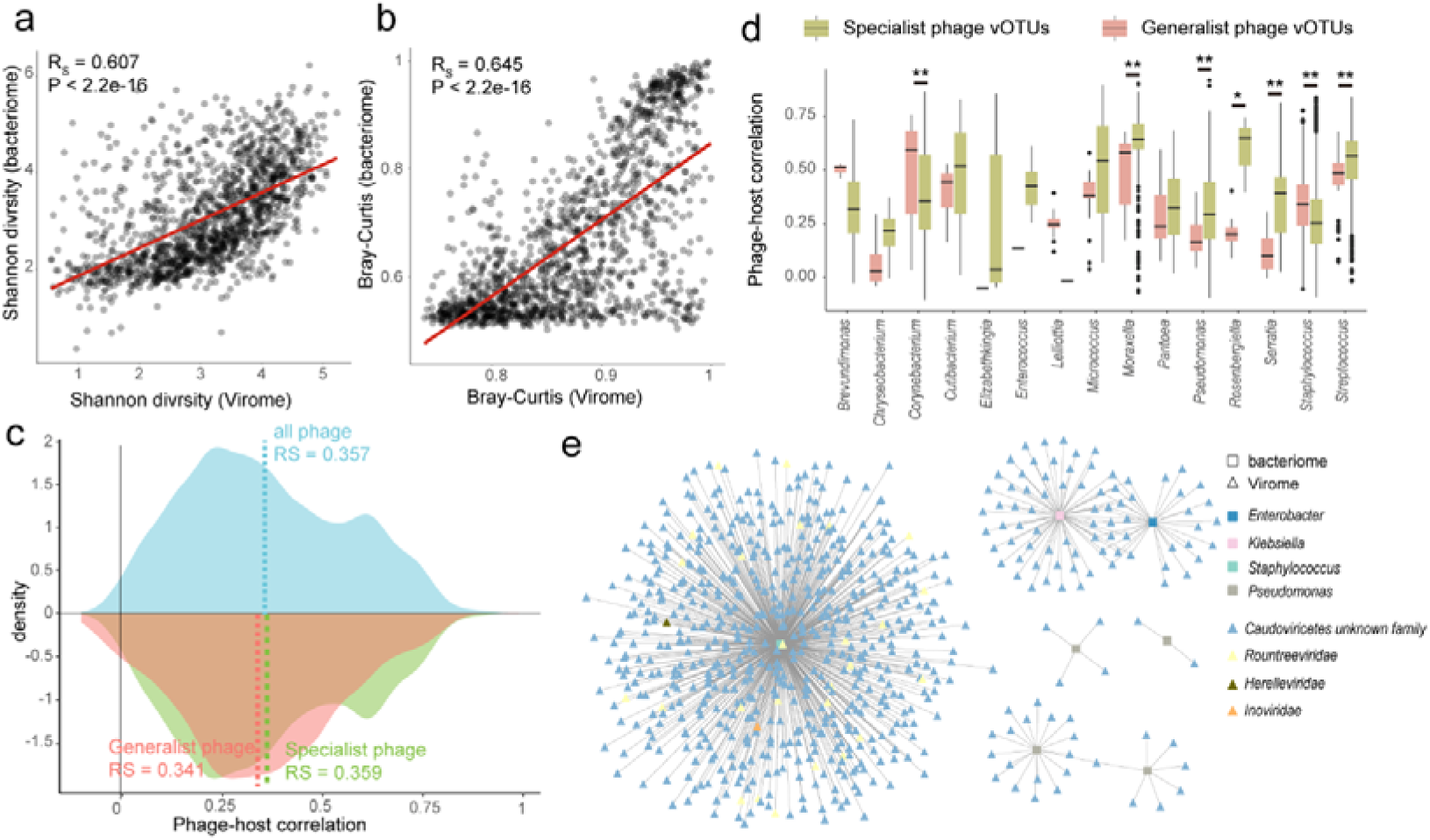
The close interactions between the skin virome and bacteriome. Comparison of α diversity (**a**) and average β diversity (**b**) between the human skin virome and bacteriome. Each circle in b represents the average Bray-Curtis distance compared with other samples. The regression line is shown in red. **c.** Distribution of phage-host correlations among 2,760 samples. Blue, green, and red respectively represent the distribution of all vOTUs, specialist phages, and generalist phages. Dashed lines indicate the average correlation within the distribution. **d.**Spearman correlation between the relative abundance of vOTUs and their predicted hosts at the genus level. In the boxplot, yellow and red represent specialist phages vOTUs and generalist phages vOTUs, respectively. **e.** Network of associations between vOTUs and their multidrug-resistant bacterial hosts. Triangles represent vOTUs, while squares represent bacteria. The calculation of correlation was performed using Spearman’s rank correlation, and the presence of a gray line indicates a significant correlation between the two variables (FDR < 0.05).

Multidrug-resistant bacteria, such as *Pseudomonas*, *Klebsiella*, and *Staphylococcus*^1, 50^, were present on the human skin. We found that numerous phages vOTUs were significantly correlated with them, particularly members of the *Caudoviricetes* including the *Rountreeviridae* and *Herelleviridae* (Figure 3e). These bacteria, of high clinical concern^51^, can cause diseases in skin, blood, lungs, gastrointestinal tract, and other parts of the body^52, 53^.

### The Biogeographic Characteristics of Viruses on the Human Skin

Human skin is characterised by different microenvironments (oily, dry and moist) and it has been shown that specific skin phenotypes are associated with distinct bacterial diversity^54-57^. In this study, three of the six datasets with samples collected from different microenvironments were analyzed. It was observed that in CNP0003934, the viral diversity (Shannon) in dry skin was significantly higher than in oily skin, similar to bacterial diversity. However, the viral diversity in oily and moist skin exhibited inconsistent trends in two dataset (the SRP002480 and SRP057859); moist skin had higher diversity than oily skin in SRP002480, while in SRP057859, oily skin exhibited higher diversity than moist skin (Figure 4a). Additionally, variations in diversity between different sites within the same skin microenvironment were observed (Figure S6 and Table S5).

**Figure 4.**
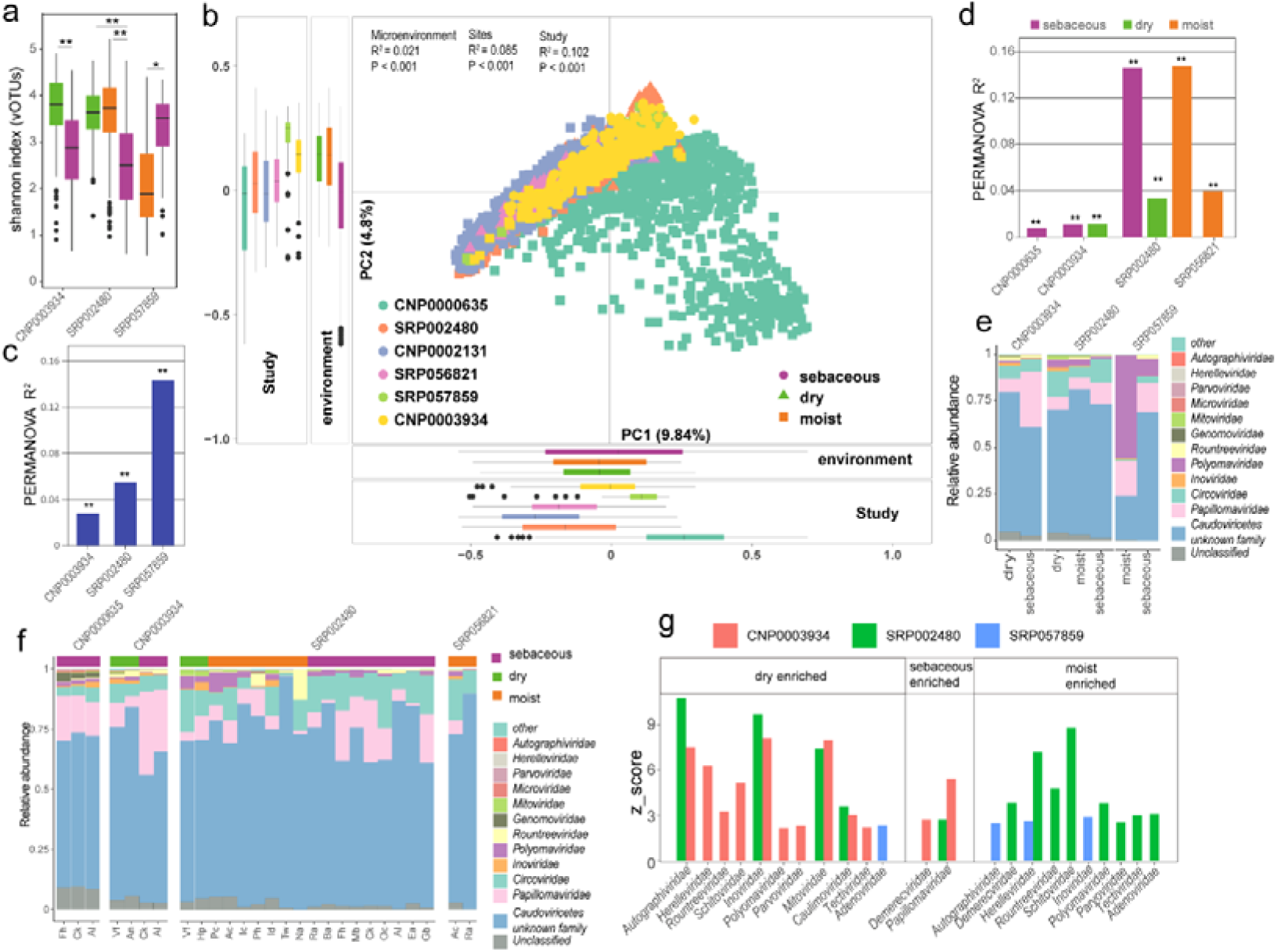
Biogeographic characteristics of the skin virome. **a.** Differences in the Shannon index at the vOTU level across different skin microenvironment within each dataset. **b.** Principal Component Analysis (PCA) based on the skin vOTU level. Different colors represent different cohorts, and shapes represent different microenvironment. Effect size (R²) and statistical significance are obtained through PERMANOVA (adonis). **c-d.** Effect size on the skin virome of different sites within the same microenvironment (c) and across various cohorts (d) and across different microenvironment. In the figure, * indicates p < 0.05, ** indicates p < 0.01. **e.** Composition at the family level of the virome in different skin microenvironment in each dataset. **f.** Composition at the family level of the virome for different sites within the same skin microenvironment in each dataset. **g.** Comparison of the relative abundance of viruses at the family level in different microenvironment within each dataset. An absolute z-score above 2 is considered statistically significant. Fh: forehead, Ck: cheek, AI: Alar crease, Ea: external auditory canal, Ra: retroarticular crease, Oc: occiput, Ba: back, Mb: manubrium, Na: nare, Ac: antecubital fossa, Id: interdigital web, Pc: popliteal fossa, Ic: inguinal crease, Vf: volar forearm, Hp: hypothenar palm, Tw: toe webspace, Tn: toenail, Ph: plantar heel, Gb: Glabella. CNP0003934: Yi_2024, CNP0000635: Zhiming_2021, SRP002480: Julia_2016, SRP056821: KernRei_2016, SRP057859: Adrian_2017, CNP0002131: Zhiming_2023.

PCoA revealed significant differences in the virome composition among the six human cohorts from different populations (PERMANOVA R² = 10.2%, p < 0.001; Figure 4b). Skin microenvironments and anatomic sites had a significant impact on the overall virome composition (PERMANOVA R² = 2.1% and R² = 8.5%, p < 0.001). Integrated data from various studies, sites, and microenvironments, we discovered that the most significant influence on the skin virome originated from the differences between studies, which accounted for 6.14% of the variation, followed by the impact of site at 3.14%, and the smallest was from the microenvironment at 0.08% (Table S6). We subsequently quantified the impact of different skin microenvironment and sites on the skin virome within each study and found that both different microenvironment and anatomical sites within the same microenvironment were significantly associated with virome composition (Figure 4cd). Furthermore, as individual heterogeneity was shown to be closely related to viral taxonomic variation, we evaluated the influence of age and sex on the skin virome across all the data. After adjusting for the effects of different studies, the impacts of age and sex were 0.08% and 0.06%, respectively (Table S6). Additionally, adjusting for the host’s age and sex did not significantly change the influence of different skin microenvironment and skin sites on the skin virome.

A compositional analysis at the viral family level across different skin microenvironments was conducted and found that unclassified vOTUs accounted for 3.21% of the viral sequences (Figure 4ef). Across all datasets, the virome of skin from different microenvironment and sites was dominated by members of the *Caudoviricetes* that have not been identified at the family level, as well as *Papillomaviridae*, *Circoviridae*, and *Inoviridae* (Figure 4ef). Oily areas were particularly enriched with the most abundant viruses on the skin, *Papillomaviridae*, while dry areas had a higher concentration of *Autographiviridae*, *Inoviridae*, *Mitoviridae* and *Retroviridae*. Moist areas were enriched with viruses from the *Herelleviridae* (Figure 4g and Table S7). Taken together, the viral composition on the skin surface exhibits microenvironment-specific and anatomic site-specific characteristics.

## Discussion

By constructing a comprehensive, non-redundant viral reference catalog from human skin samples across diverse microenvironments and geographic locations, our study has addressed several gaps in the current understanding of the skin virome. We discovered a significant number of vOTUs that do not match any known sequences, constituting approximately 90.85% (n = 2,610) of our findings. This suggests that viral diversity is more abundant and varied than previously acknowledged. Our analysis also sheds light on the intricate relationships these viruses maintain with their host microenvironments and the broader skin microbiome, revealing how specific microenvironmental and anatomical sites influence viral roles and adaptations.

A major challenge in our study stemmed from the metagenomic sequencing data, which, unlike more targeted approaches, did not undergo a VLP enrichment step nor included a host depletion step. The absence of VLP enrichment meant that our samples were not specifically concentrated for viruses, thereby including a substantial amount of non-viral genetic material. This lack of enrichment complicates the detection and characterization of viral entities, particularly those that are less abundant or that exist in complex biological matrices where they are overshadowed by more dominant microbial forms. Furthermore, the lack of a host depletion step prior to sequencing resulted in a high background noise of human DNA^58, 59^. This background significantly complicates the bioinformatic processing required to distinguish viral DNA from the human DNA, which is predominant in skin samples^5, 6^. Consequently, identifying and cataloging viral sequences required the implementation of stringent filtering criteria to ensure the accuracy and reliability of our viral sequence identifications.

In our research, we used multiple methods to detect viral sequences, including:

1. DeepVirFinder: A deep learning-based tool designed for predicting viral sequences.
2. VIBRANT: A machine learning-based method for identifying viral sequences.
3. geNomad: Utilizes gene content and deep neural networks to recognize viral sequences.

Additionally, we performed alignments against the complete human genome to eliminate sequences that may be of human origin. Finally, we used CheckV to evaluate and select viral sequences of medium quality or higher. This multi-faceted approach enhances the accuracy and reliability of our viral detection and characterization in metagenomic samples.

Despite these efforts, the limitations inherent in our approach underscore the need for more refined methodologies in future studies. Employing VLP enrichment and host DNA depletion in the initial stages of sample preparation would likely allow for a more comprehensive and detailed exploration of the skin virome. These steps would not only increase the yield of viral sequences but also improve the signal-to-noise ratio, facilitating the detection of rare and novel viruses that could be crucial for understanding the full scope of viral impact on skin health and disease.

Additionally, our results highlight the impact of microenvironmental factors and sites on the composition of the virome.. The variation in viral communities across different microenvironment and regions illustrates how these viruses have adapted to specific local microenvironments. This specificity to anatomical sites and microenvironment indicates that localized treatment approaches may be crucial, as the skin virome appears to be closely adapted to its surrounding conditions.

## Conclusion

This extensive exploration of the human skin virome provides a deeper understanding of the viral components of the skin microbiome and highlights their potential role in influencing skin health and disease. As we continue to unravel the complex interactions within the skin microbiome, these findings not only pave the way for innovative therapeutic approaches, but also enhance our understanding of skin biology. The way forward is to integrate these findings into broader biological and clinical research to fully realise the potential of the skin virome in promoting health and treating disease.

## Acknowledgements

This work was supported by the Ministry of Science and Technology of China (2022ZD0211600 and 2023YFC3603300), CAMS Innovation Fund for Medical Sciences (2019-I2M-5-066), the National Natural Science Foundation of China (82341108) and Innovation Team Fund from Guangzhou Municipal Science and Technology Bureau (2024D03J0015). The authors thank China National GeneBank and Guangdong Provincial Key Laboratory of Genome Read and Write (No. 2017B030301011), Shenzhen Key Laboratory of Neurogenomics (BGI Genomics, No. CXB201108250094A) for support in sequencing and analysis.

## Author contributions

J.X. and Z.L. conceived the study. Z.L. performed the analyses. J.X., J.W., C.N. and J.K. supervised the work and provided funding. Z.L., S.L. C.H.,Y.C., H.Z.,Y.S., J.K. and J.X. wrote the paper. Y.Z., Y.C. reviewed and revised the manuscript critically. All authors read, edited and approved the paper.

